# Low-input ATAC&mRNA-Seq: a simple and robust method for simultaneous dual-omics profiling with low cell number

**DOI:** 10.1101/2021.02.25.432879

**Authors:** Ruifang Li, Sara A Grimm, Paul A Wade

## Abstract

Deciphering epigenetic regulation of gene expression requires measuring the epigenome and transcriptome jointly. Single-cell multi-omics technologies have been developed for concurrent profiling of chromatin accessibility and gene expression. However, multi-omics profiling of low-input bulk samples remains challenging. Therefore, we developed low-input ATAC&mRNA-seq, a simple and robust method for studying the role of chromatin structure in gene regulation in a single experiment with thousands of cells, to maximize insights from limited input material by obtaining ATAC-seq and mRNA-seq data simultaneously from the same cells with data quality comparable to conventional mono-omics assays. Integrative data analysis revealed similar strong association between promoter accessibility and gene expression using the data of low-input ATAC&mRNA-seq as using single-assay data, underscoring the accuracy and reliability of our dual-omics assay to generate both data types simultaneously with just thousands of cells. We envision our method to be widely applied in many biological disciplines with limited materials.

## Introduction

Joint profiling of the epigenome and transcriptome is needed to unravel epigenetic regulation of gene expression. Conventional high-throughput transcriptome profiling and epigenome mapping technologies, such as MNase-seq (Schones, Cui et al. 2008), DNase-seq (Boyle, Davis et al. 2008), and FAIRE-seq (Giresi, Kim et al. 2007), typically require large amounts of input material (i.e. millions of cells or more); therefore, they are not suitable for many cutting-edge studies requiring transcriptome and epigenome analyses of very small amounts of input material. Assay for transposase-accessible chromatin using sequencing (ATAC-seq) (Buenrostro, Giresi et al. 2013) is a state-of-the-art low-input technology widely used for studying chromatin structure. Combining ATAC-seq with RNA-seq offers a powerful approach to understand the role of chromatin structure in regulating gene expression. Recently, several single-cell multi-omics profiling assays have been developed for concurrent measurement of chromatin accessibility and gene expression in the same cell (Cao, Cusanovich et al. 2018, Liu, Liu et al. 2019, Ma, Zhang et al. 2020, Xing, Farran et al. 2020). These methods hold considerable promise for low-input material, but they cannot be easily adopted in regular biological laboratories because of high cost and complex workflows. In addition, the sparse and noisy nature of single-cell data remains a limitation.

It remains a challenge to profile the epigenome and transcriptome simultaneously with a limited amount of material in many biological disciplines. Therefore, we developed low-input ATAC&mRNA-seq, a simple, rapid and low-cost method for fast simultaneous profiling of chromatin accessibility and gene expression with a small number of cells. As a proof-of-concept, we applied low-input ATAC&mRNA-seq to E14 mouse embryonic stem cells (mESCs) using 5K, 10K, and 20K cells as input, and compared the resulting ATAC-seq and mRNA-seq data with conventional single-assay data. We found that our method generated comparable high-quality data even with just 5K cells, thus providing a unique solution to multi-omics profiling with limited material.

## Results

We developed low-input ATAC&mRNA-seq (Figure 1) to simultaneously profile chromatin accessibility and gene expression with a small number of cells. Instead of isolating the nuclei as in Omni-ATAC (Corces, Trevino et al. 2017), our method uses a one-step membrane permeabilization and transposition of whole cell for ATAC-seq. As for mRNA sequencing, our method uses direct mRNA isolation from the cell lysate with Dynabeads^®^ Oligo (dT)_25_ followed by solid-phase cDNA synthesis, and then employs Tn5 transposase to directly fragment and tag mRNA/cDNA hybrids to form an amplifiable library for sequencing. Our mRNA-seq strategy enables a seamless on-bead process in one tube, which not only simplifies sample handling but also minimizes sample loss. With the greatly simplified workflow, our method takes only ∼ 4 hours from cells to sequencing ready ATAC-seq and mRNA-seq libraries, dramatically reducing hands-on time.

**Figure 1.**
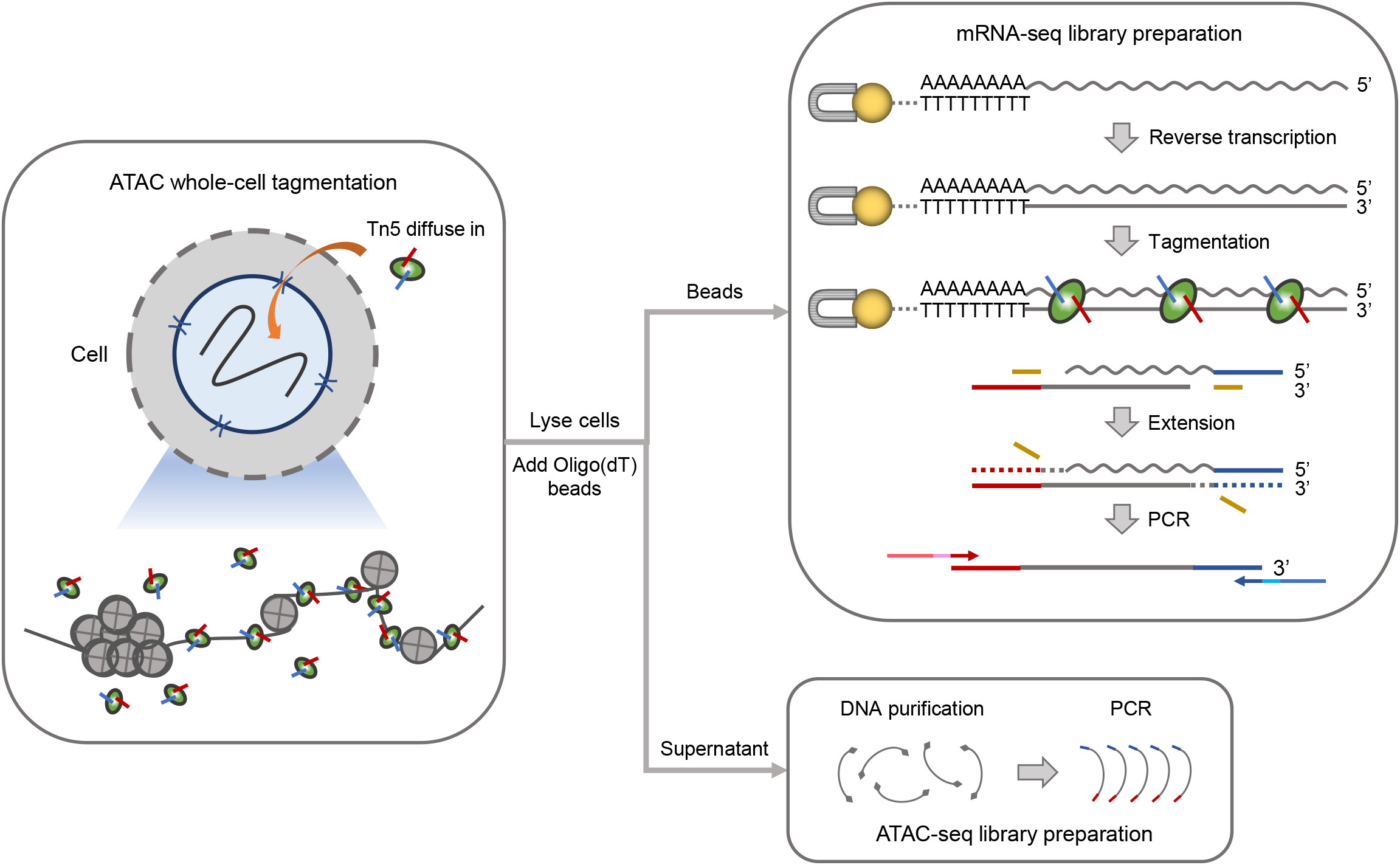
A schematic overview of low-input ATAC&mRNA-seq workflow. Harvested cells were washed and then permeabilized with mild detergent (indicated by holes in cell membrane) to facilitate the entry of Tn5 into the nuclei to tagment open chromatin regions. Tagmented cells were then lysed and Dynabeads Oligo (dT)_25_ were added into the cell lysate to capture mRNA. After magnetic separation, tagmented genomic DNA in the supernatant was purified and further amplified with indexed PCR to construct ATAC-seq library, while mRNA captured on beads was reverse transcribed using the bead-bound oligo (dT) as primer. The mRNA/cDNA hybrids were then directly tagmented by Tn5, and after initial end extension, the tagmented cDNA was amplified with indexed PCR to prepare mRNA-seq library. Wavy and straight gray lines represent RNA and DNA, respectively. Dotted lines represent the extended fragment.

To benchmark our method, we performed low-input ATAC&mRNA-seq on E14 mESCs with low input cell numbers (5K, 10K, and 20K) and compared the resulting data with single-assay data from Omni-ATAC-seq and conventional bulk mRNA-seq to assess data quality. Omni-ATAC (Corces, Trevino et al. 2017) is an improved protocol for ATAC-seq with dramatically reduced mitochondrial reads and enhanced signal-to-noise ratio. Even though our method used whole cell instead of isolated nuclei for transposition reaction, it achieved similarly low percentage of mitochondrial reads contamination (<10%) comparable to Omni-ATAC (Figure 2A). Notably, the duplication rate was low across our ATAC-seq libraries and it increased only marginally with decrease in the number of input cells (Figure 2A), indicating high complexity of the libraries even when starting with just 5K cells. Our ATAC-seq libraries exhibited the characteristic nucleosome periodicity in fragment size distribution along with transcription start site (TSS) enrichment (Figure 2B and Figure S1), which are typical of a successful ATAC-seq experiment. The patterns were consistent across our ATAC-seq libraries with different input cell numbers, albeit with slight differences in the TSS enrichment score and visibility of the nucleosome periodicity (Figure S1). Overall, our ATAC-seq libraries were of comparable quality to Omni-ATAC libraries in terms of ATAC-seq quality control (QC) metrics.

**Figure 2.**
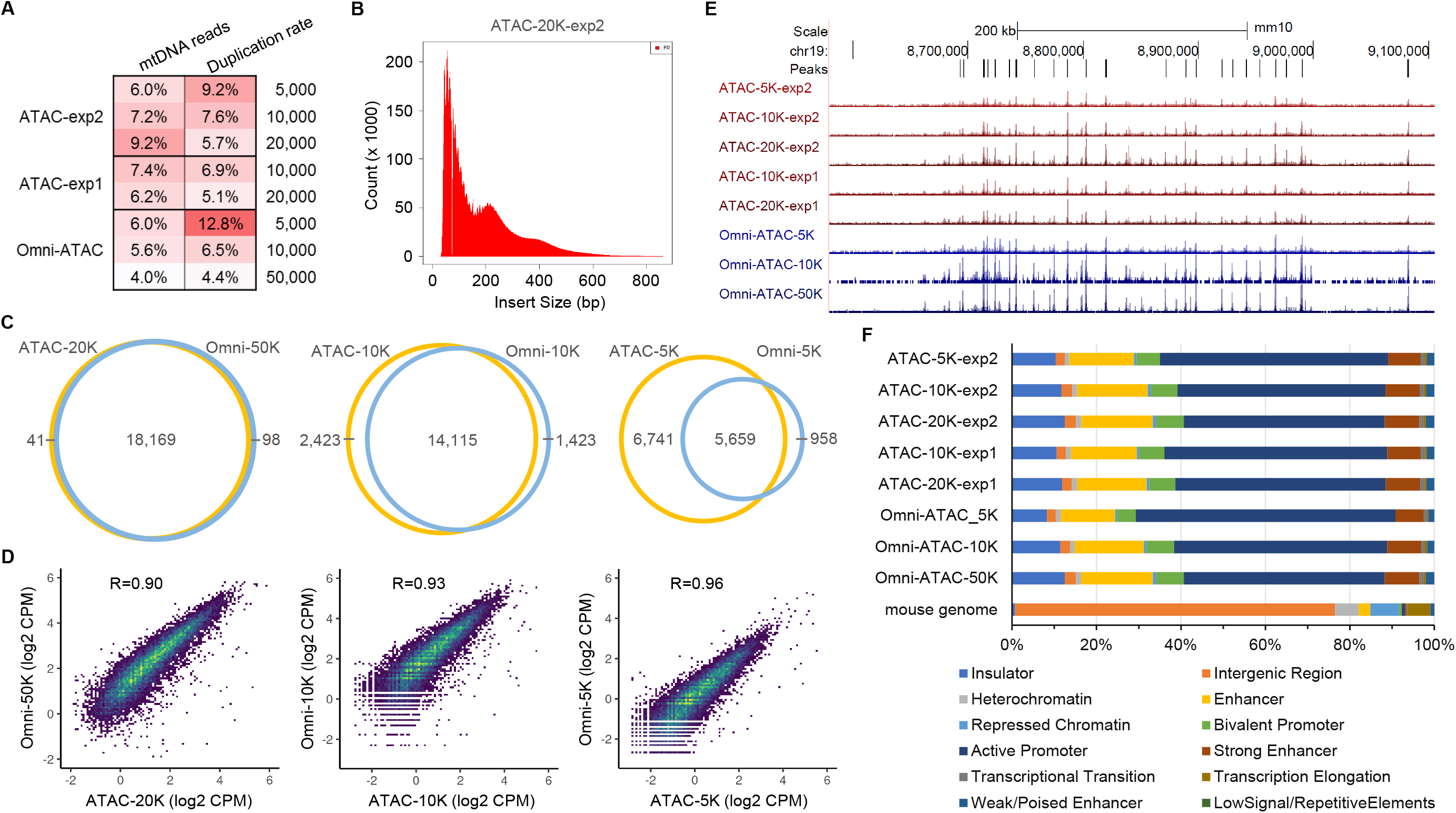
Comprehensive evaluation of ATAC-seq data of low-input ATAC&mRNA-seq compared with Omni-ATAC-seq. Two independent low-input ATAC&mRNA-seq experiments (exp1 and exp2) as well as Omni-ATAC-seq were performed on E14 mESCs with different input cell numbers. (A) Heatmap representation of ATAC-seq quality control metrics including duplication rate and the percentage of reads mapped to mitochondrial DNA (mtDNA). Lighter color is used to depict the more desirable value of each metric, along with the numeric values shown for each sample. The numbers of input cells used in each sample were shown on the right of the heatmap. (B) Fragment size distribution of a representative ATAC-seq library prepared with low-input ATAC&mRNA-seq. (C) Venn diagrams showing overlap of ATAC-seq peaks identified by low-input ATAC&mRNA-seq and Omni-ATAC-seq. (D) Density scatter plots displaying correlation of ATAC-seq data generated with low-input ATAC&mRNA-seq and Omni-ATAC-seq. Each dot represents an individual peak in the unified peak set with viridis color scale indicating density. Pearson’s *r* value was shown at the top of each plot. (E) UCSC genome browser view of ATAC-seq coverage tracks at chr19:8,579,587-9,105,586. (F) Bar graph showing the proportion of ATAC-seq peaks and the mouse genome falling into each chromatin state of mESC.

Next, we compared the identified accessible chromatin regions (ATAC-seq peaks) by the two methods. Compared with standard Omni-ATAC using 50K cells as input, our method obtained a similar number of peaks with only 20K cells (Omni-50K: 18,267; ATAC-20K: 18,210); while with 10K and 5K cells as input, our method detected more peak regions than Omni-ATAC (Figure 2C), suggesting that our method requires even fewer cells than Omni-ATAC, which is largely attributable to minimized sample loss with the one-step approach in our simplified ATAC workflow. Importantly, the majority of ATAC-seq peaks identified with our method overlapped with Omni-ATAC peaks (Figure 2C) and ATAC-seq signals at unified peak regions were highly correlated between our method and Omni-ATAC (Pearson’s *r* ≥ 0.90) over a range of input cell numbers (Figure 2D), indicating remarkably high consistency in enrichment between our ATAC-seq data and Omni-ATAC-seq data as exemplified in the coverage tracks (Figure 2E). Open chromatin regions encompass several key features of the epigenome, including active and poised regulatory regions. To verify that our ATAC-seq data correctly identified those regulatory features, we investigated the epigenomic contexts of ATAC-seq peaks with respect to chromatin states of mESC as defined by ChromHMM model (Pintacuda, Wei et al. 2017). We observed similar distribution of ATAC-seq peaks identified by our method and Omni-ATAC (Figure 2F), with ∼51% of peaks located in ‘Active Promoter’, ∼6% in ‘Bivalent Promoter’, ∼8% in ‘Strong Enhancer’, ∼16% in ‘Enhancer’, ∼2% in ‘Weak/Poised Enhancer’, and ∼11% in ‘Insulator’, consistent with previous findings that ATAC-seq peaks predominantly overlap with active and poised chromatin states while barely overlap with repressed and inactive chromatin states (Tarbell and Liu 2019). Taken together, the comprehensive analyses demonstrated that our method is comparable to Omni-ATAC in generating high-quality ATAC-seq data to identify the key regulatory regions controlling cell identity.

To evaluate the quality of our mRNA-seq data, we compared it with previously published E14-mESC mRNA-seq data (Ramisch, Heinrich et al. 2019) generated using Illumina TruSeq Stranded mRNA kit, which is the prevailing library preparation kit for conventional bulk mRNA-seq. Compared with TruSeq libraries, our mRNA-seq libraries showed even lower percentage of rRNA contamination (Avg. 0.06% vs 0.37%) (Figure 3A) and exhibited similar read distribution across known gene features with ∼ 82% of reads mapped to exons, ∼ 10% to introns, and ∼ 8% to intergenic regions (Figure 3B), validating the mRNA origin of the libraries and negligible genomic DNA contamination with direct mRNA isolation from the cell lysate. Inspection of read coverage over gene bodies revealed a bias towards the 3′-end of genes (3′ bias) in our mRNA-seq data (Figure 3C and 3D; Figure S2). Nevertheless, in total 12,438 genes were detected in our mRNA-seq data with a minimum expression threshold of 0.5 TPM. The number of detected genes was slightly lower in our mRNA-seq data than in TruSeq data (12,438 vs 14,114), which was expected given that the input cell numbers were two orders of magnitude less in our experiment than in TruSeq experiment. Importantly, 93.25% of the expressed genes (11,599 out of 12,438) were concordantly detected in TruSeq data (Figure 3E); moreover, the measured gene expression levels exhibited strong correlation between our method and TruSeq method across the transcriptome (Spearman’s *ρ* = 0.90, Figure 3F), suggesting that our method performs comparably to conventional bulk mRNA-seq in terms of gene expression measurement but requires substantially fewer cells.

**Figure 3.**
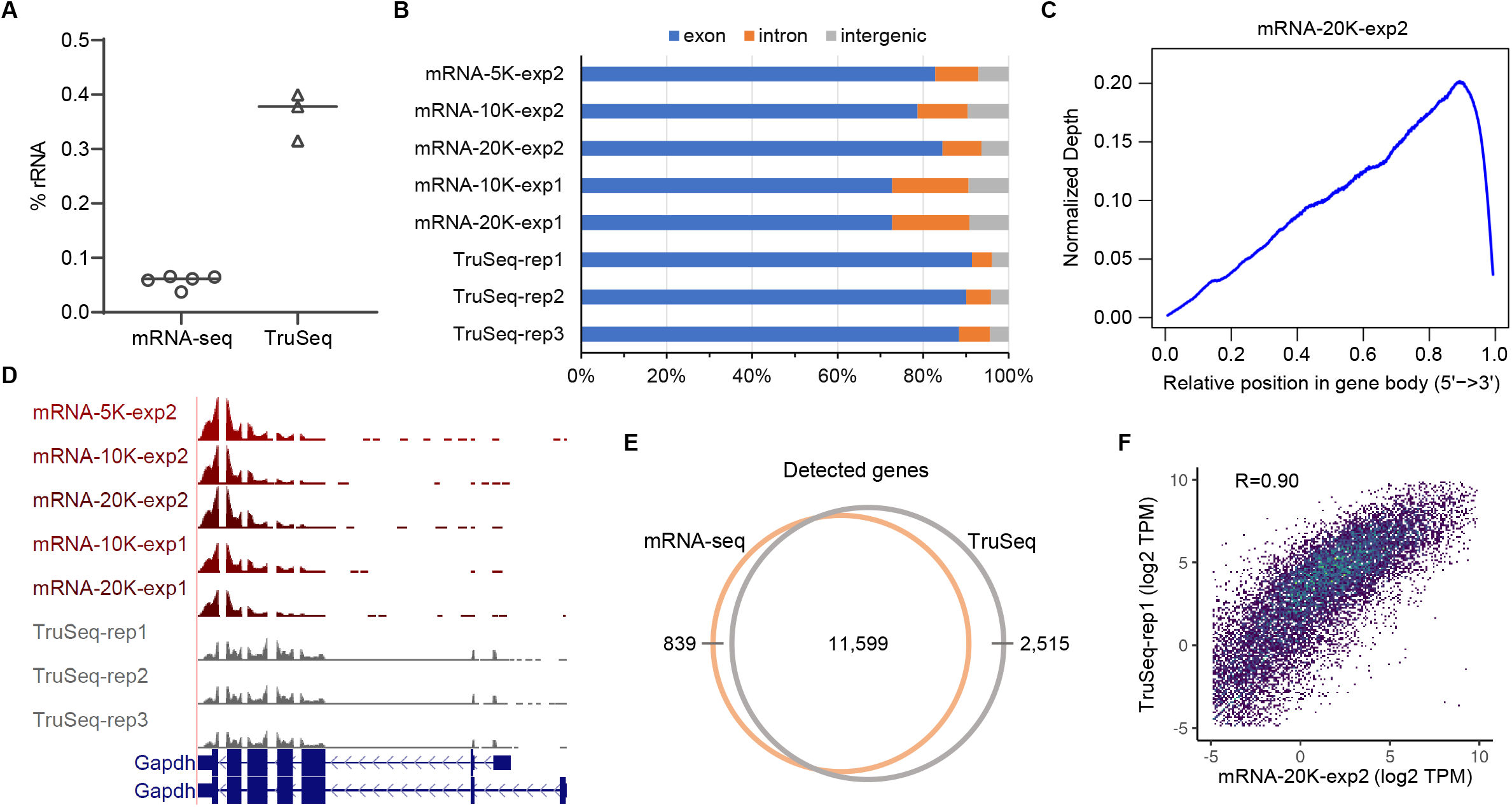
Comprehensive evaluation of mRNA-seq data of low-input ATAC&mRNA-seq compared with conventional bulk mRNA-seq. (A) Column scatter graph showing the percentage of reads mapped to rRNA in each sample with the line indicating the median of each group. (B) Distribution of mapped reads across known gene features (exons, introns, and intergenic regions). (C) Gene body coverage profile of a representative mRNA-seq library prepared with low-input ATAC&mRNA-seq. The curve was smoothed over 15 consecutive points in the plot. (D) UCSC genome browser view of mRNA-seq coverage tracks at Gapdh gene locus. (E) Venn diagram showing overlap of detected genes (TPM ≥ 0.5) in low-input ATAC&mRNA-seq and conventional bulk mRNA-seq. (F) Density scatter plot displaying correlation of gene expression measured by low-input ATAC&mRNA-seq and conventional bulk mRNA-seq. Each dot represents a gene with viridis color scale indicating density. Spearman’s *ρ* value was shown at the top of the plot.

mRNA sequencing reveals the transcriptional state of genes whereas chromatin accessibility mapping with ATAC-seq uncovers the associated regulatory landscape. With joint profiling of accessible chromatin and mRNA expression in the same cells, our method would enable a direct link of transcriptional regulation to its output. Indeed, integrative analysis of promoter accessibility and gene expression showed that ‘open’ promoters were correlated with relatively high gene expression while ‘closed’ promoters were associated with low gene expression (Figure 4A), which was similar to that observed by integrating conventional single-assay mRNA-seq and ATAC-seq data generated with substantially more input cells (Figure 4B), suggesting that our dual-omics profiling method performed comparably well in integrative data analysis. Notably, consistent results were obtained across different input cell numbers (Figure 4A), demonstrating the reliability of our method to detect association of chromatin accessibility and gene expression with just thousands of cells.

**Figure 4.**
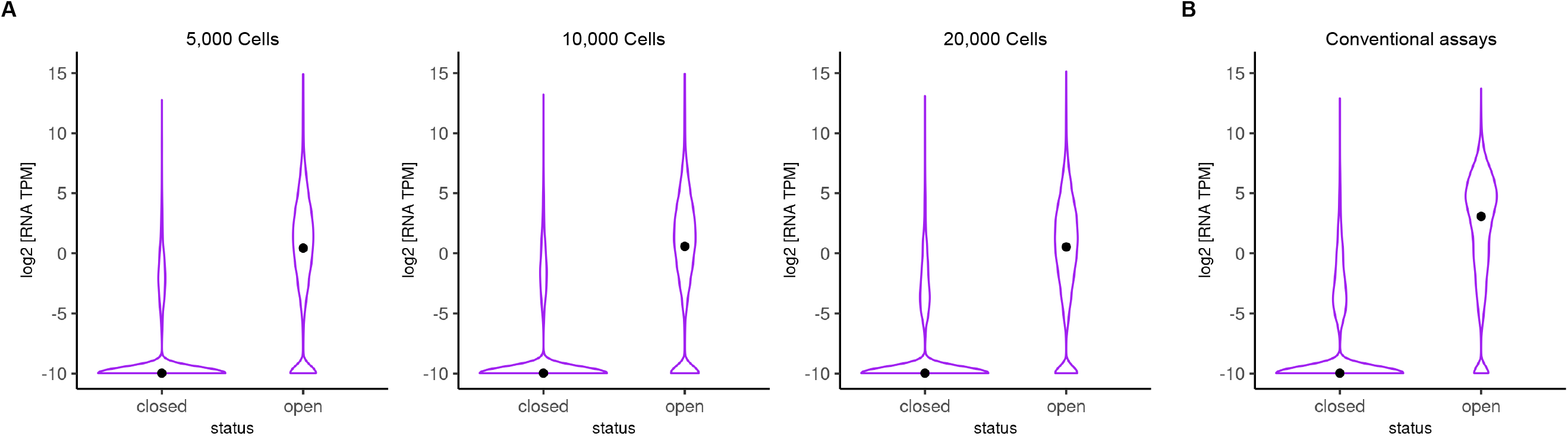
Integrative analysis of promoter accessibility and gene expression. Violin plots showing mRNA expression levels of genes with ‘open’ versus ‘closed’ promoter by integrating ATAC-seq and mRNA-seq data generated with (A) low-input ATAC&mRNA-seq and (B) conventional mono-omics assays (Omni-ATAC and TruSeq mRNA sequencing) using substantially larger numbers of input cells in the same context (publicly available data). Black points indicate the median of mRNA expression levels in each category. For plotting purpose, a floor value of 0.001 TPM was applied.

Lastly, to demonstrate the robustness of our method, we compared samples from the same batch but using different numbers of input cells. Remarkably, ATAC-seq peaks identified in those samples largely overlapped (Figure 5A and Figure S4A), and ATAC-seq signals at peak regions showed very high correlation in pairwise comparisons of those samples (Pearson’s *r* = 0.96-0.99) (Figure 5B and Figure S4B), indicating high consistency in ATAC-seq enrichment despite different input amount. Likewise, a similar number (∼ 11,000) of genes were detected across the mRNA-seq samples regardless of input cell numbers, and the vast majority (∼ 90%) of the detected genes were shared among the mRNA-seq samples (Figure 5C and Figure S4C). Moreover, the measured gene expression levels were highly consistent across the samples with different input cell numbers (Pearson’s *r* > 0.99) (Figure 5D and Figure S4D). Collectively, these analyses showed the robustness of our method against variation in the number of input cells. Next, to assess reproducibility of the data generated with our method, we compared samples from two independent experiments performed on different days. Irrespective of the number of input cells (10K or 20K), the independent samples consistently exhibited large overlap of ATAC-seq peaks (Figure 6A) and high correlation of ATAC-seq signals at unified peaks (Pearson’s *r* ≥ 0.98) (Figure 6B); similarly, they also showed high concordance in mRNA-seq data with ∼ 92.5% of the detected genes in common (Figure 6C) and highly consistent gene expression levels (Pearson’s *r* ≥ 0.99) (Figure 6D), altogether indicating high reproducibility of the ATAC-seq and mRNA-seq data generated with our method. In conclusion, our method is robust and reproducible with low cell input to obtain both data types simultaneously.

**Figure 5.**
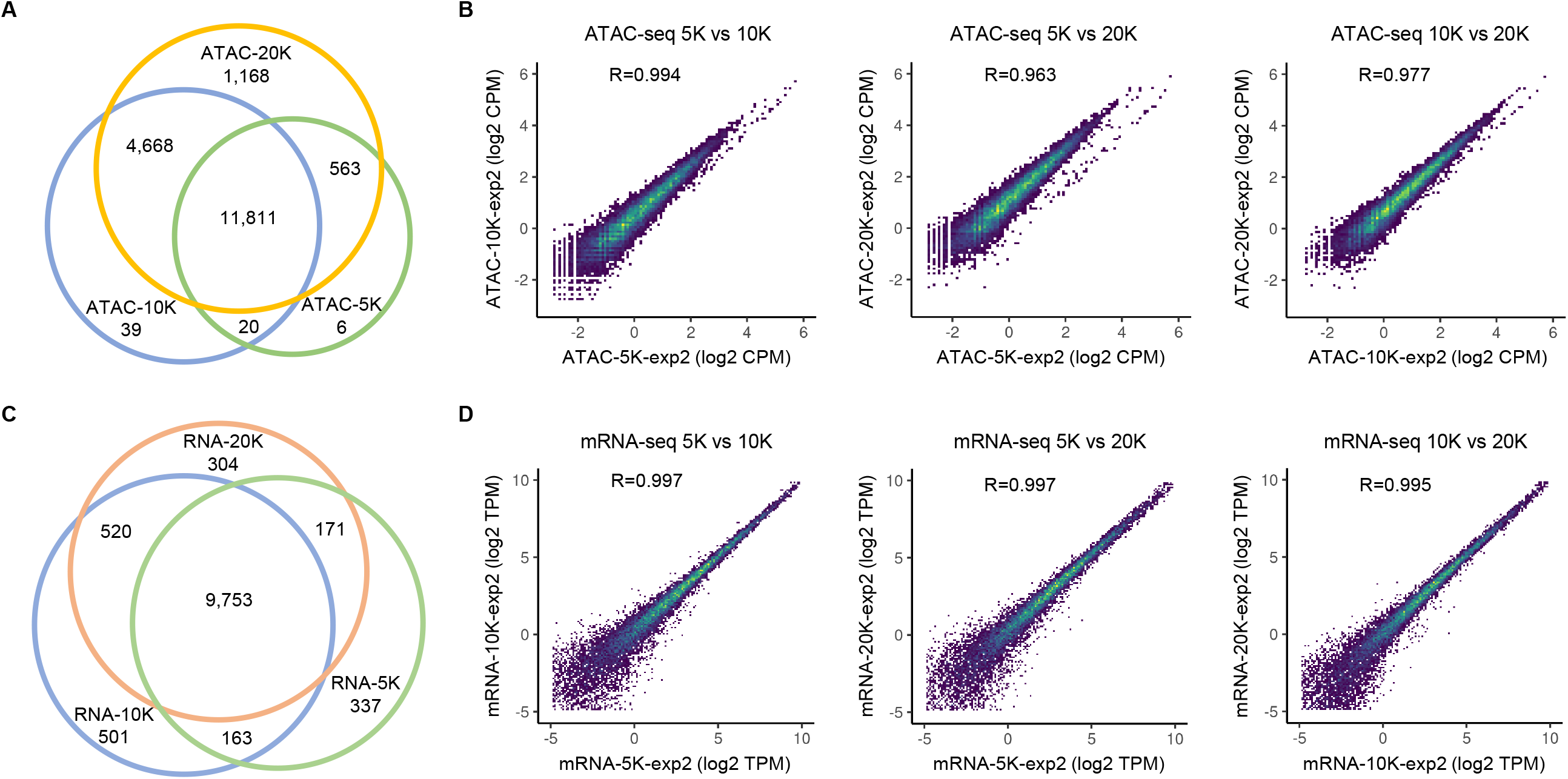
Evaluating robustness of low-input ATAC&mRNA-seq against variation in the number of input cells. (A) Venn diagram showing overlap of ATAC-seq peaks identified with 20K, 10K, and 5K input cells. (B) Density scatter plots displaying pairwise correlations of ATAC-seq data generated with 20K, 10K, and 5K input cells. Each dot represents an individual peak in the unified peak set with viridis color scale indicating density. Pearson’s *r* value was shown at the top of each plot. (C) Venn diagram showing overlap of detected genes (TPM ≥ 0.5) with 20K, 10K, and 5K input cells. (D) Density scatter plots displaying pairwise correlations of gene expression measured with 20K, 10K, and 5K input cells. Each dot represents a gene with viridis color scale indicating density. Pearson’s *r* value was shown at the top of each plot.

**Figure 6.**
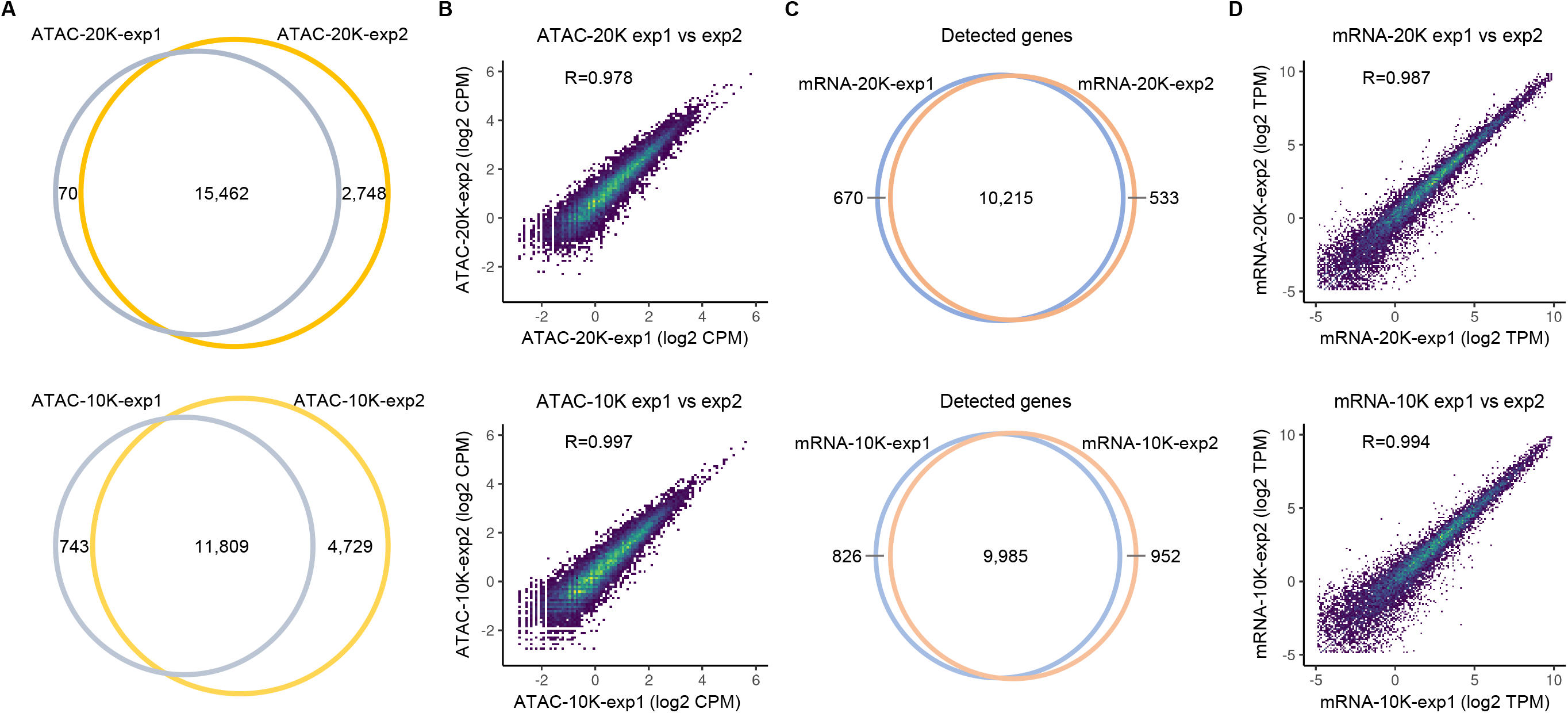
Assessing reproducibility of data generated with low-input ATAC&mRNA-seq. (A) Venn diagrams showing overlap of ATAC-seq peaks identified in two independent experiments using the same numbers of input cells. (B) Density scatter plots displaying correlation of ATAC-seq data of the two independent experiments. Each dot represents an individual peak in the unified peak set with viridis color scale indicating density. Pearson’s *r* value was shown at the top of each plot. (C) Venn diagrams showing overlap of detected genes (TPM ≥ 0.5) in the two independent experiments. (D) Density scatter plots displaying correlation of gene expression measured in the two independent experiments. Each dot represents a gene with viridis color scale indicating density. Pearson’s *r* value was shown at the top of each plot.

## Discussion

We invented a dual-omics profiling method (low-input ATAC&mRNA-seq) to simultaneously measure chromatin accessibility and mRNA expression in the same cells with low cell number. Although multi-omics profiling of low-input material is challenging, we succeeded in that by improving both ATAC-seq and mRNA-seq procedures to minimize sample loss during the process. Our method uses whole cell instead of nuclei for chromatin tagmentation in ATAC (Figure 1). By removing the nuclei isolation step, it not only simplifies the experimental procedure but also avoids cell loss without significantly affecting the quality of ATAC-seq data. As a result, our method requires even fewer cells than Omni-ATAC to identify more accessible chromatin regions encompassing key regulatory features in the epigenome, including active and poised promoters, enhancers, and insulators (Figure 2). The advantage became more noticeable especially when starting with much fewer cells, as demonstrated by the superior performance of our method with 5K cells in terms of ATAC-seq library complexity, peak number, and consistency with larger-input samples (Figure 2). In addition, the improvement of ATAC procedure enables downstream mRNA-seq to measure both cytoplasmic and nuclear transcripts. With whole-cell transcriptome profiling, it greatly increases the amount of mRNA available for library preparation and ensures fair comparison with publicly available mRNA-seq datasets as most use RNA isolated from whole cells.

Conventional bulk mRNA-seq library construction involves many laborious and time-consuming steps, including RNA extraction, mRNA purification with poly-A selection or ribosomal RNA depletion, mRNA fragmentation, reverse transcription, second-strand cDNA synthesis, end repair, adaptor ligation, and PCR amplification (Mortazavi, Williams et al. 2008, Cui, Lin et al. 2010), with purification procedures needed between the enzymatic steps causing inevitable sample loss. To minimize sample loss during mRNA-seq library preparation for low-cell-input samples, we replaced the complex traditional method with a simple ‘one-tube’ method that consists of only three seamless steps including direct mRNA isolation from cell lysate with Dynabeads^®^ Oligo (dT)_25_, on-bead cDNA synthesis, tagmentation of mRNA/cDNA hybrids and PCR amplification (Figure 1). The whole procedure was performed on beads, enabling simple and rapid wash and buffer change between the steps without requiring any laborious purification. Our mRNA-seq approach greatly simplifies the workflow and minimizes sample loss. With just thousands of input cells, it generated acceptable mRNA-seq data that was comparable to conventional bulk mRNA-seq data in terms of read distribution across gene features, number of detected genes, and gene expression levels, albeit with inferior coverage uniformity over gene body (Figure 3). The 3′ bias in gene body coverage is a common phenomenon also observed in other similar approaches such as SHERRY (Di, Fu et al. 2020) and TRACE-seq (Lu, Dong et al. 2020), two recently published RNA-seq methods similar to our mRNA-seq approach in using oligo(dT) primed cDNA synthesis and Tn5-mediated direct tagmentation of RNA/cDNA hybrids for library preparation.

In summary, by coupling the simplified ATAC procedure with the novel mRNA-seq approach, low-input ATAC&mRNA-seq can simultaneously profile both chromatin accessibility and gene expression with low cell numbers ranging from 5K to 20 K cells. The ATAC-seq and mRNA-seq data generated with our method were highly reproducible and strongly correlated with counterpart data of conventional mono-omics methods (Figure 2-6), demonstrating the accuracy and reliability of our method to generate both data types simultaneously with just thousands of cells. Our method uses commercially available off-the-shelf reagents and requires only basic molecular biology equipment. The speed, low-input requirement, technical simplicity and robustness of our method make it appealing, especially in situations when it is challenging to collect large numbers of cells. Hence, we envision our method to be widely useful in many biological disciplines with limited materials. Furthermore, as a scalable method, low-input ATAC&mRNA-seq holds promise for even fewer input cells with further optimization of the protocol, which would also reduce reaction costs by using proportionally less amounts of reagents.

## Methods

### Low-input ATAC&mRNA-seq

Harvested E14 mouse ESCs were washed twice with cold PBS and then aliquoted into three Eppendorf tubes with 20K, 10K, and 5K cells, respectively. The cells were then spun down at 500 x g, 4 °C for 5 min in a pre-chilled fixed-angle microcentrifuge. The supernatant was removed carefully without disturbing the cell pellet. The cell pellets were then resuspended in 20 μl, 10 μl, and 8 μl of transposition mix (25 μl 2×TD buffer, 2.5 μl Tn5 (Illumina), 16.5 μl PBS (Invitrogen), 0.5 μl 1% digitonin (Promega), 0.5 μl 10% Tween-20 (Roche), 1 μl RNase inhibitor (Invitrogen), and 4 μl nuclease-free water), respectively, by pipetting up and down six times. Permeabilization/transposition reactions were incubated at 37 °C for 30 min in a thermomixer with shaking at 1000 rpm. Once the incubation finished, EDTA and LiCl were added to a final concentration of 10 mM and 0.5 M, respectively, followed by adding 100 μl of Lysis/Binding Buffer in Dynabeads^®^ mRNA DIRECT™ Micro Kit (Invitrogen) and pipetting up and down to lyse the cells. After complete cell lysis, 20 μl pre-washed Dynabeads^®^ Oligo (dT)25 were added into the cell lysate and the mixture was incubated at room temperature for 5 min with continuous rotation to allow the mRNA to anneal to the oligo(dT) on Dynabeads. The sample tubes were then placed on a Dynal magnet for 1 minute to separate the mRNA and genomic DNA (gDNA). The supernatant containing tagmented gDNA was transferred into a new Eppendorf tube followed by DNA purification with MinElute PCR purification kit (Qiagen). Meanwhile, the mRNA captured on beads was washed extensively according to the manual of Dynabeads^®^ mRNA DIRECT™ Micro Kit (Invitrogen). The Dynabeads-mRNA complex was then resuspended in 20 μl of reverse transcription reaction mix without any primer prepared using SuperScript™ IV First-Strand Synthesis System (Invitrogen), and the reaction was incubated initially at 50°C for 5 minutes and then at 55°C for 10 minutes. The bead-bound oligo(dT) was used as primer for first strand cDNA synthesis, and as a result, the mRNA/cDNA hybrids were covalently bound to the Dynabeads. Once reverse transcription was completed, the PCR tubes were immediately placed on the magnet for 30 seconds and the supernatant was then removed. The Dynabeads-mRNA/cDNA complex was washed twice in 100 μl of ice-cold 10 mM Tris-HCl and then resuspended in 5 μl of ice-cold 10 mM Tris-HCl. Direct tagmentation of the mRNA/cDNA hybrids and PCR amplification were performed on beads using Nextera XT DNA Library Prep Kit (Illumina); the Reference Guide was followed exactly for 20K and 10K samples, while the amounts of reagents used for 5K sample were scaled down by half. In the meantime, previously purified ATAC-DNA was amplified using NEBNext^®^ High-Fidelity 2X PCR Master Mix (NEB) with 10 cycles of PCR. The ATAC-seq and mRNA-seq libraries were purified with AMPure XP beads (Beckman Coulter) and then quantified with Qubit dsDNA HS Assay Kit (Invitrogen).The libraries were pooled and sequenced on Illumina NextSeq550 with paired-end 75-bp reads.

### Omni-ATAC-seq

Omni-ATAC-seq was performed on E14 mESCs following Omni-ATAC protocol (Corces, Trevino et al. 2017) with slight modifications. Briefly, harvested mESCs were counted and then aliquoted into three Eppendorf tubes with 50K, 10K, and 5K cells, respectively. The cells were washed once with cold ATAC-seq resuspension buffer (RSB; 10 mM Tris-HCl pH 7.4, 10 mM NaCl, and 3 mM MgCl_2_ in water) followed by lysing the cells on ice for 3 min in 50 μl of ATAC-seq RSB containing 0.1% NP40 (Roche), 0.1% Tween-20 (Roche), and 0.01% Digitonin (Promega). After cell lysis, 1 ml of cold ATAC-seq RSB containing 0.1% Tween-20 (without NP40 or digitonin) was added and then mixed by inverting the tubes. Isolated nuclei were spun down at 500 x g, 4 °C for 10 min in a pre-chilled fixed-angle centrifuge. After removing the supernatant, the nuclei of 50K, 10K, and 5K samples were resuspended in 50 μl, 10 μl, and 5 μl of transposition mix (25 μl 2x TD buffer, 2.5 μl Tn5 (Illumina), 16.5 μl PBS (Invitrogen), 0.5 μl 1% digitonin (Promega), 0.5 μl 10% Tween-20 (Roche), 5 μl H2O), respectively, by pipetting up and down six times. Transposition reactions were incubated at 37 °C for 30 min in a thermomixer with shaking at 1000 rpm. The reactions were cleaned up with MinElute PCR Purification Kit (Qiagen), and the purified DNA was amplified using NEBNext^®^ High-Fidelity 2X PCR Master Mix (NEB) with 10 cycles of PCR. The ATAC-seq libraries were purified with AMPure XP beads (Beckman Coulter) and quantified with Qubit dsDNA HS Assay Kit (Invitrogen). The libraries were then pooled and sequenced on Illumina NextSeq550 with paired-end 35-bp reads.

### Publicly available data used in this work

Previously published datasets of conventional mRNA-seq and ATAC-seq on E14 mESCs were downloaded as raw fastq files from GEO series GSE120376 (GSM3399470, GSM3399471, GSM3399472, GSM3399494) (Ramisch, Heinrich et al. 2019).

### ATAC-seq data analysis

ATAC-seq fastq files were filtered to remove all entries with a mean base quality score below 20 for either read in the read pair. Adapters were removed via Cutadapt v1.12 (Martin 2011) with parameters “-a CTGTCTCTTATA −O 5 -q 0”, and the trimmed reads were further filtered to exclude those with length less than 30 bp. The remaining filtered and trimmed read pairs were mapped against the mm10 reference assembly via Bowtie2 v2.1.0 (Langmead and Salzberg 2012) with parameters “-X 2000 --fr --end-to-end --very-sensitive”, followed by filtering with samtools v1.3.1 (Li, Handsaker et al. 2009) at MAPQ5. Reads mapped to chrM were ignored in all downstream analysis. Duplicate mapped read pairs were removed by Picard tools MarkDuplicates.jar (v1.110) (http://broadinstitute.github.io/picard). Only the 9 bp at the 5’ end of each read was retained for downstream analysis. Coverage tracks for genome browser views were generated using BEDtools v2.24.0 (Quinlan and Hall 2010) genomeCoverageBed with depth normalized to 10 million read ends per sample and then converted to bigWig format with UCSC utility bedGraphToBigWig (http://hgdownload.soe.ucsc.edu/admin/exe/).

### Peak calling and peak overlap among samples

MACS2 v2.1.1 (Zhang, Liu et al. 2008) was used for initial peak calls per sample with parameters “callpeak -g mm -q 0.0001 --keep-dup=all --nomodel --extsize 9”, followed by merging peaks within 200bp with BEDtools v2.24.0 mergeBed. To facilitate comparisons across samples, a single set of unified peaks was generated by collapsing called peaks and identifying only peak regions that overlap from at least 3 of 8 samples via BEDtools v2.24.0 unionBedGraphs, again followed by merging peaks within 200bp with BEDtools v2.24.0 mergeBed. Unified peaks less than 50 bp in width were discarded, as were peaks outside canonical chromosomes (chr1-19, X, Y). Analysis of shared peaks among samples was performed using subsets of unified peaks that overlap the MACS2 peak calls of given samples. Briefly, the unified peak set was intersected individually with MACS2 peak calls from each sample in comparison. The subset of unified peaks in the intersection represented the total peaks in the sample. These subsets of unified peaks were then intersected with each other to determine the overlap of peaks among samples.

### Overlap of ATAC-seq peaks with chromatin states

Chromatin states of mESC as defined by ChromHMM (Pintacuda, Wei et al. 2017) were downloaded from https://github.com/guifengwei/ChromHMM_mESC_mm10. To investigate the epigenomic contexts of ATAC-seq peaks, each peak was assigned to the chromatin state with which it showed the most overlap.

### Promoter accessibility stratification

Promoter was defined as the region +/- 1Kb of the annotated TSS. To determine promoter accessibility, ATAC-seq signal was quantified by counting number of 9-mer read ends per promoter region with BEDtools v2.24.0 multiBamCov and then converted to CPM (counts per million). Promoters were stratified as either “open” or “closed” with a hard cutoff of 1.45 CPM, which was derived from the bimodal distribution of ATAC-seq signals at promoters (Figure S3). For genes with multiple TSS, we selected the one with the maximum ATAC-seq CPM that was averaged over all evaluated samples.

### RNA-seq data analysis

RNA-seq fastq files were filtered to remove all entries with a mean base quality score below 20 for either read in the read pair. Filtered read pairs were mapped against the mm10 reference assembly via STAR v2.5 (Dobin, Davis et al. 2013) with parameters “--outSAMattrIHstart 0 -- outFilterType BySJout --alignSJoverhangMin 8 --limitBAMsortRAM 55000000000 -- outSAMstrandField intronMotif --outFilterIntronMotifs RemoveNoncanonical”. Coverage tracks were generated with STAR v2.5 with parameters “--runMode inputAlignmentsFromBAM -- outWigType bedGraph --outWigStrand Unstranded --outWigNorm RPM”, followed by conversion to bigWig format via UCSC utility bedGraphToBigWig.

Read counts per gene were determined via featureCounts (Subread v1.5.0-p1) (Liao, Smyth et al. 2014) with parameters “-s0 -Sfr -p” and then converted to TPM (transcripts per million). The gene models used in this study were from NCBI RefSeq annotations limited to only curated transcripts, with a GTF format version downloaded from the UCSC Genome Browser (http://hgdownload.soe.ucsc.edu/goldenPath/mm10/bigZips/genes/, dated January 10, 2020).

Gene body coverage profile was generated by calculating the number of mapped reads that overlap with genomic bins tiled over exonic regions of a given gene model (where each bin covers 0.1%) via BEDtools v2.24.0 intersectBed, then aggregating over all gene models, and finally normalizing by total counts.

## Data availability

The data described in this manuscript have been deposited in the NCBI Gene Expression Omnibus (GEO) with accession number GSE165478.

## Acknowledgements

We thank Dr. Hua Wang in Helin lab for providing the E14 mESCs. This work was supported by donations to the CER from The Metropoulos Family Foundation and The Ambrose Monell Foundation.

## Figure legends

**Figure S1.**
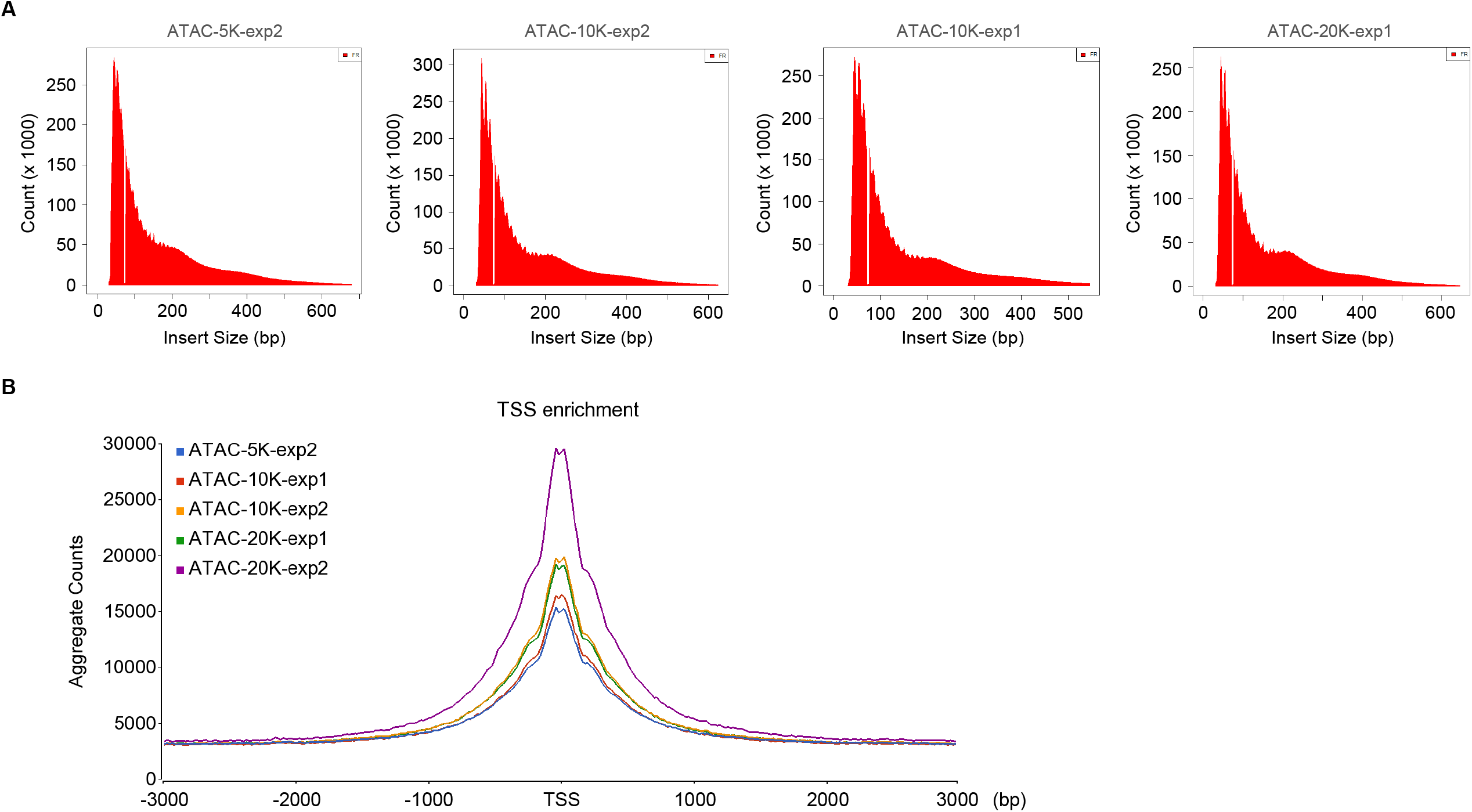
(A) Fragment size distribution and (B) TSS enrichment of ATAC-seq libraries prepared with low-input ATAC&mRNA-seq using different numbers of input cells.

**Figure S2.**
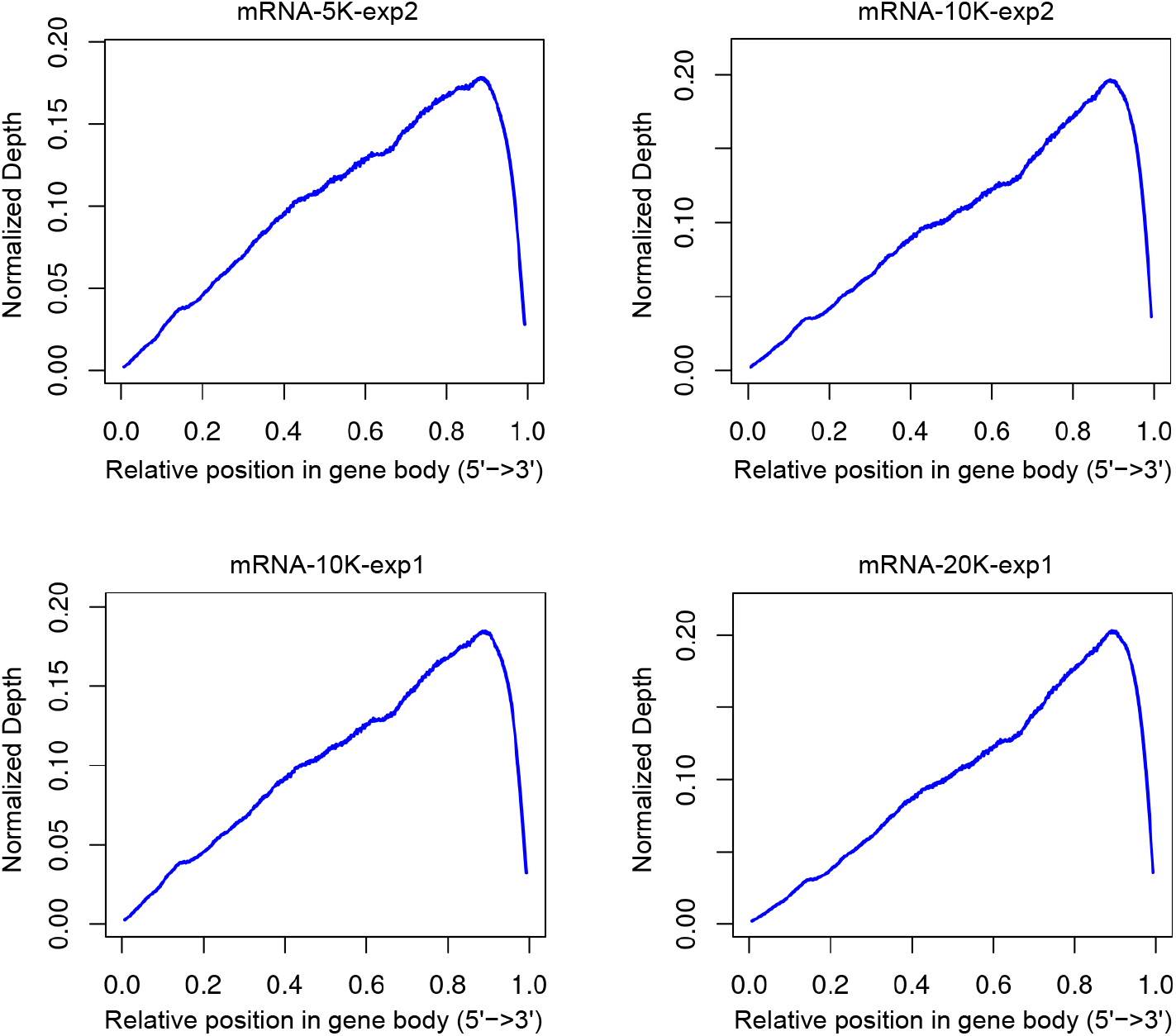
Gene body coverage profiles of mRNA-seq libraries prepared with low-input ATAC&mRNA-seq.

**Figure S3.**
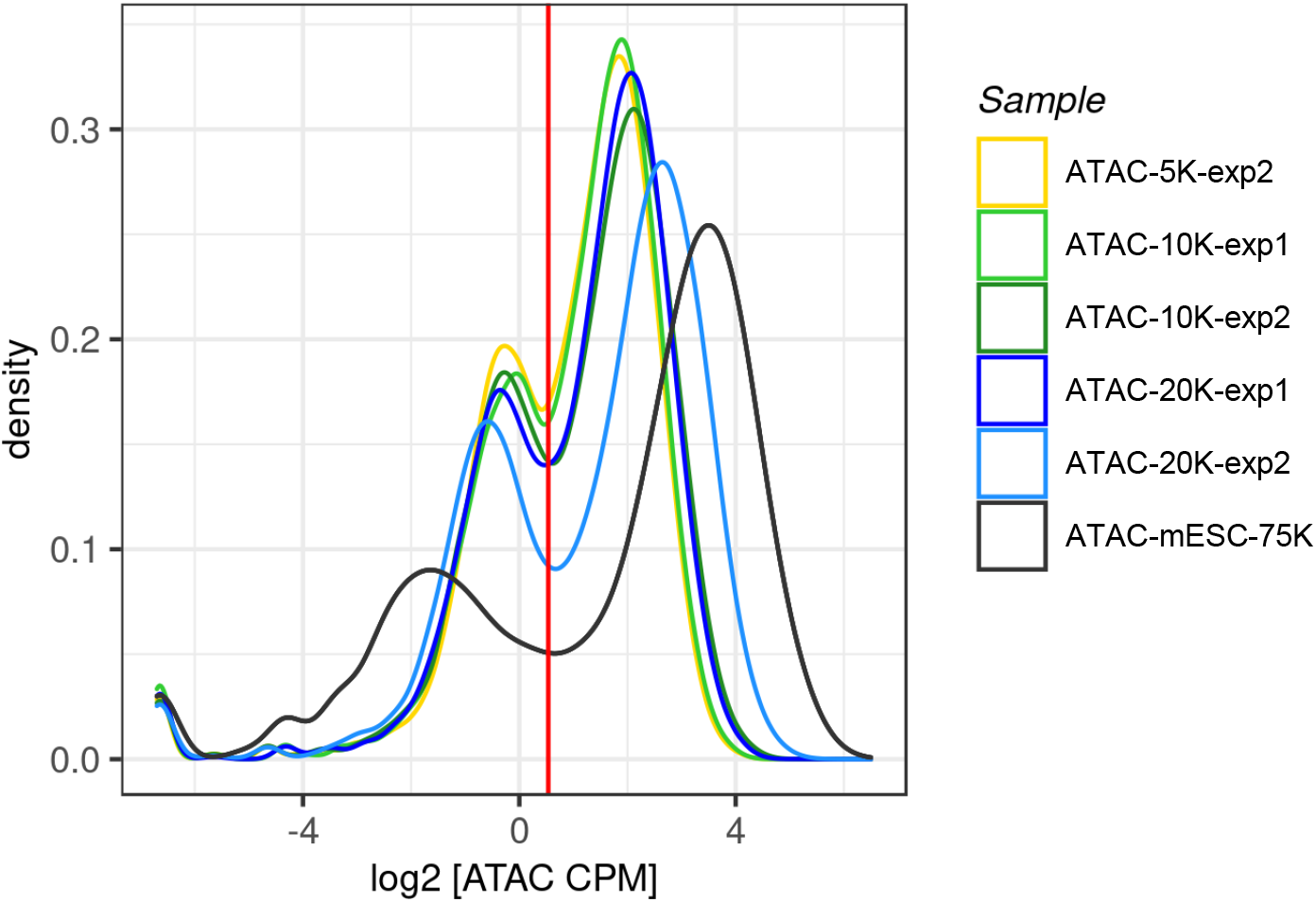
Density plot showing the distribution of ATAC-seq signals at promoter regions in each ATAC-seq sample. The red line indicates the selected cutoff for stratifying ‘closed’ and ‘open’ promoters.

**Figure S4.**
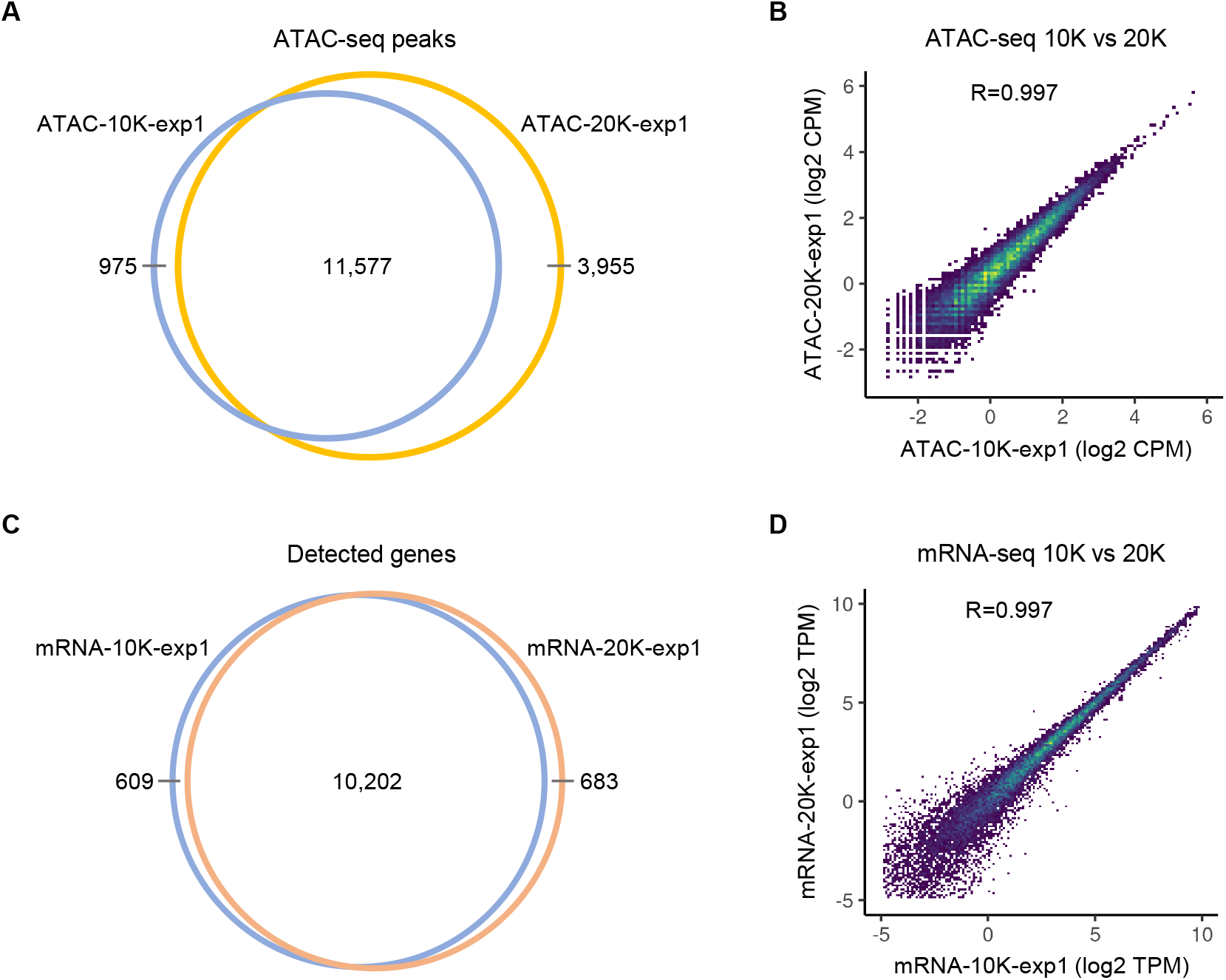
Evaluating robustness of low-input ATAC&mRNA-seq against variation in the number of input cells. Data was shown for an independent experiment (exp1) performed on a different day from data shown in Figure 5. (A) Venn diagram showing overlap of ATAC-seq peaks identified with 20K and 10K input cells. (B) Density scatter plot displaying correlation of ATAC-seq data generated with 20K and 10K input cells. Each dot represents an individual peak in the unified peak set with viridis color scale indicating density. Pearson’s *r* value was shown at the top of the plot. (C) Venn diagram showing overlap of detected genes (TPM ≥ 0.5) with 20K and 10K input cells. (D) Density scatter plot displaying correlation of gene expression measured with 20K and 10K input cells. Each dot represents a gene with viridis color scale indicating density. Pearson’s *r* value was shown at the top of the plot.

